# Joint Representation of Color and Shape in Convolutional Neural Networks: A Stimulus-rich Network Perspective

**DOI:** 10.1101/2020.08.11.246223

**Authors:** JohnMark Taylor, Yaoda Xu

## Abstract

To interact with real-world objects, any effective visual system must jointly code the unique features defining each object. Despite decades of neuroscience research, we still lack a firm grasp on how the primate brain binds visual features. Here we apply a novel network-based stimulus-rich representational similarity approach to study color and shape binding in five convolutional neural networks (CNNs) with varying architecture, depth, and presence/absence of recurrent processing. All CNNs showed near-orthogonal color and shape processing in early layers, but increasingly interactive feature coding in higher layers, with this effect being much stronger for networks trained for object classification than untrained networks. These results characterize for the first time how multiple visual features are coded together in CNNs. The approach developed here can be easily implemented to characterize whether a similar coding scheme may serve as a viable solution to the binding problem in the primate brain.

## Introduction

Natural visual experience comprises a juxtaposition of different visual features, such as an object’s shape, color, position, size, and orientation. To recognize an object under different viewing conditions, our visual system must successively reformat and “untangle” the different features to make object identity information explicitly available to a linear readout process in a manner that is tolerant to variations in other features, an ability that has been hailed as the hallmark of primate high-level vision (DiCarlo & Cox, 2007; Hong et al., 2016).

Meanwhile, our interaction with the world often involves objects with uniquely defined features, such as grabbing the blue pen on the desk. How would an object representation that sheds all its identity-irrelevant features support our ability to interact with specific objects? One possibility is that different visual features are initially processed separately and are bound together via attention (i.e., Feature Integration Theory, Treisman & Gelade, 1980). Despite decades of neuroscience research, the coding mechanism for such a binding process remains unknown, with existing proposals facing various challenges. For example, Singer (1999) proposed that neurons coding for different features of the same object could engage in synchronous oscillations, serving as a binding signal, but it is unclear how such a signal would be generated and read out (Shadlen & Movshon, 1999). Alternatively, there might exist neurons that encode particular feature conjunctions; however, this view collides with the problem of “combinatorial explosion”: there are more possible feature conjunctions than neurons in the brain. The space of possible mechanisms requires further exploration.

Recently, convolutional neural networks (CNNs) have achieved human-level object recognition performance (Kriegeskorte, 2015; Yamins & Dicarlo, 2016; Rajalingham, et al., 2018; Serre, 2019). Specifically, these CNNs have been trained to disregard identity-irrelevant object features to correctly identify objects across different viewing conditions, thereby forming transformation-tolerant visual object representations much like those in high-level primate vision. However, despite their success in object recognition, CNNs largely remain “black boxes”, with details of their internal processing poorly understood (e.g., Serre, 2019). Several studies have examined how individual features are encoded in CNNs, with some finding that coding for object identity-irrelevant features increases in higher CNN layers (Hong et al., 2016), and others reporting the color encoding characteristics of CNN units (Flachot & Gegenfurtner, 2018; Rafegas & Vanrell, 2018). However, to our knowledge no study has examined how CNNs encode *combinations* of features during the course of information processing. Because CNNs are not trained to interact with specific objects but simply to produce the correct object labels at the end of its processing, it is possible that different features are initially encoded in an entangled, intermingled fashion, and are gradually separated, with object identity information made explicit over the course of processing (DiCarlo & Cox, 2007). Alternatively, CNN architecture and training for object recognition may *automatically* give rise to interactive coding of object features in later stages of processing, without needing a separate binding operation. This could constitute a novel binding mechanism that has not been considered before in neuroscience research. Thus, studying how CNNs jointly encode different object features during the course of visual information processing is not only timely in its own right, but also provides us with a unique opportunity to gain insight into the potential computational algorithm that a successful object recognition system may use to code different object features together. Equally importantly, given that the internal representations of CNNs are fully image computable and freely inspectable, CNNs provide ideal testing grounds for developing analysis methods to study feature coding across an entire processing hierarchy and with a large number of objects, and generating hypotheses that can be tested in biological visual systems.

In this study, we examined how an object’s color and shape may be coded together during visual processing in CNNs. We employ a network-based, stimulus-rich approach in which we characterize the joint representations of these two features not for a few pairs of objects at a few processing stages as is traditionally done in neuroscience and vision research, but rather, across a large number of objects and across the entire processing hierarchy of a CNN. With this approach, we found that coding for color and shape becomes increasingly interactive throughout CNN processing. These results characterize for the first time how multiple visual features are coded together in CNNs. The approach developed here can be easily implemented to characterize whether the primate brain may use a similar coding scheme to solve the binding problem.

## Results

In this study, we examined in detail how color and naturalistic object shape features may be represented together in five CNNs trained for object recognition using ImageNet (Deng et al., 2009) images. These CNNs, chosen for their high object recognition performance, architectural diversity, and prevalence in the literature, included AlexNet (Krizhevsky, Sutskever, & Hinton, 2012), VGG19 (Simonyan & Zisserman, 2015), GoogLeNet (Szegedy et al., 2015), ResNet-50 (He, Zhang, Ren, & Sun, 2015), and CORNet-S (Kubilius et al., 2018). Specifically, AlexNet was included for its high object recognition performance, relative simplicity, and prevalence in the literature. VGG19, GoogLeNet and ResNet-50 were chosen based on their high object recognition performance and architectural diversity. Both AlexNet and VGG19 have a shallower network structure, whereas GoogLeNet and ResNet-50 have a deeper network structure. CORNet-S is a shallow recurrent CNN designed to approximate the structure of the primate ventral visual pathway, and exhibits high correlation with neural and behavioral metrics. This CNN has recently been argued to be the current best model of the primate ventral visual regions (Kar et al., 2019). We sampled between 6 to 9 layers in each of these CNNs (Table 1).

**Table 1.** The five CNNs included in the present study and the layers sampled in each CNN.

| **Network**<br>(Layers Used/Total Layers) | **Layers Used** |
| --- | --- |
| AlexNet (8/25) | Conv1, Conv2, Conv3, Conv4, Conv5, FC1, FC2, FC3 |
| CORnet-S (7/42) | Conv1, Relu2, Relu5, Relu8, Relu11, AvgPool1, FC1 |
| GoogLeNet (6/144) | Conv1, MaxPool2, MaxPool5, MaxPool11, AvgPool1, FC1 |
| ResNet-50 (6/177) | Conv1, Relu4, Relu8, Relu14, AvgPool1, FC1 |
| VGG19 (9/47) | Conv1, MaxPool1, MaxPool2, MaxPool3, MaxPool4, MaxPool5, FC1, FC2, FC3 |

We used representational similarity analysis (RSA, Kriegeskorte & Kievit, 2013) to characterize how color and shape information is represented together in these networks. For most analyses, we studied a set of 50 objects (chosen from a set created by Brady et al., 2013), each colored in 12 colors calibrated in CIELUV color space (Figure 1a). Two versions of each object were shown: a textured version, with internal object detail preserved, and a silhouette version, comprising a global shape contour without internal details (Figure 1b).

**Figure 1.**
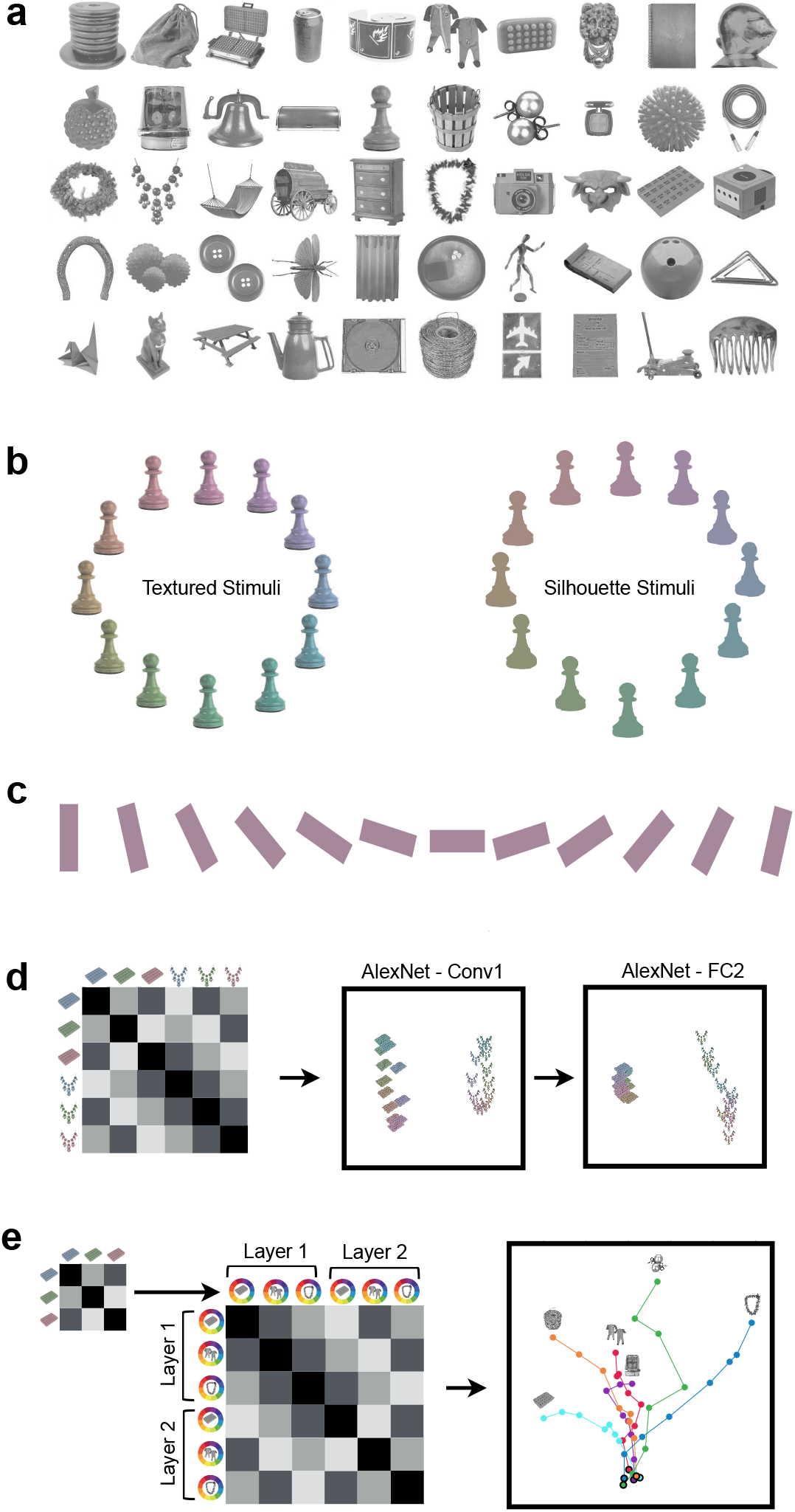
Stimuli used and example color space characterization using RSA and MDS. a. The 50 objects included in the main stimulus set, chosen from an initial set of 500 objects to maximize their mean pairwise pattern similarity in AlexNet FC2. b. The 12 isoluminant and iso-saturated colors (based on the CIELUV color space) and the two versions of the object shapes used in the main analysis. Objects appeared either with their original textures preserved, or as uniformly shaded “silhouette” stimuli. c. The 12 oriented bar stimuli used in a control analysis. d. An illustrative color similarity matrix (left) and actual MDS plots showing the representational structure of two example objects each in the 12 colors calibrated in CIELUV color space from Conv1 and FC2 of AlexNet (right). Pairwise correlations were first obtained from these two objects in the 12 colors (3 colors were illustrated here) to construct a color similarity matrix for each layer. The first two dimensions of this similarity matrix were then projected onto the 2D space using MDS. While the similarity spaces of these objects have a similar elliptical pattern at the beginning of AlexNet, by the end of processing the color spaces of these objects are substantially different. e. An illustrative color space similarity matrix (left) and an actual MDS plot showing the color spaces of six example objects over the course of processing in AlexNet (right). Color spaces were computed separately for each of these objects in each sampled layer of AlexNet (example color space depicted by the small matrix on the left), and the resulting color spaces (only three objects and two layers illustrated here) were correlated with one another to construct a color space similarity matrix (note this is a second order correlation matrix, different from the color similarity matrix illustrated in d). The first two dimensions of this similarity matrix were then projected onto the 2D space using MDS. Each dot in the MDS plot represents the color space of a given object at a given layer and each trajectory traces the color space of a given object. The dot corresponding to the initial layer has a black outline, and the dot corresponding to the final layer is marked by a picture of the object for that trajectory. While the color spaces of different objects are initially very similar, by the end of processing they have substantially diverged.

All of our analyses involved first computing the *color space* of a given object that captures how similarly the different colors of that object are coded at a particular CNN layer. Subsequent analyses then involved (1) comparing the color spaces across objects within each layer, to determine whether color and shape are encoded independently versus interactively in that layer, and examining how this is affected by variations in stimuli, analysis parameters, and network training regime; (2) comparing the color space for each object across layers, to determine how color information for each object is transformed over the course of processing; (3) examining whether color space differences across objects are preserved across layers, and (4) whether the shape similarity of two objects predicts their color space similarity. To our knowledge, these analyses provide the first in-depth and comprehensive network-based description of how colors and shapes are coded together in CNNs.

### Visualizing Color Space Representation Across Objects and CNN Layers

As our primary analysis, we applied RSA to examine the extent to which coding for color varies across object shapes, and the extent to which the magnitude of this variability changes across CNN layers. As an initial exploratory analysis, we visualized how the color spaces of two example objects may differ at the beginning and end of processing in AlexNet (Figure 1d). Specifically, we extracted the activation patterns for the 12 colors of these two objects (textured versions) from the first and the penultimate layers of AlexNet (Conv1 and FC2). Within each layer, we performed all pairwise Pearson correlation among the 24 patterns to create a representational similarity matrix (RSM). Using multidimensional scaling (MDS), we visualized the resulting representational similarity space projected onto 2D space, with a closer distance between a pair of colored objects indicating more similar representations (Figure 1d). For these two objects, color was coded similarly at the beginning of the network, as reflected by the similar elliptical shape of the color spaces; by contrast, the color spaces of these two objects were substantially different by the end of processing.

To generalize from these two objects and examine how the color space for different objects might diverge over layers, we visualized the evolution of the color spaces of six example objects over the course of processing in AlexNet (Figure 1e). To do this, for each object (textured versions) and for each sampled layer of AlexNet, we first constructed a “color space” RSM by performing all pairwise Pearson correlations of the patterns associated with the 12 different colors of that object. We vectorized the off-diagonal values of this RSM to create a “color space” vector. Next, we performed all pairwise correlations of these “color space” vectors across objects and layers to form a “color space similarity” RSM that quantifies how similarly color is coded in different objects and layers. We then used MDS to visualize the resulting representational similarity space (Figure 1e). In these objects, color was initially coded in a very similar manner (as reflected by the dense clustering of the bold-outlined dots representing the different color spaces in the initial layers of processing), but color coding increasingly diverged as processing proceeded in the network (as reflected by the separation of the dots at the end of processing, indicated by the object icons next to the dots).

### Quantifying Color Space Differences Across Objects within a CNN Layer

To quantify the color space divergence among different objects within a layer and over the course of processing, we computed the averaged pairwise color space vector correlations for the 50 objects in each layer of each CNN, and for both the textured and silhouette stimuli. We further quantified using regression analysis whether this mean between-object color space similarity significantly declines over the course of processing, and whether this decline varies significantly between the textured and silhouette versions of the stimuli (see Methods for analysis details). For the entire set of 50 objects, several patterns of results, shown in Figure 2a, could be possible: the color spaces of different objects might be highly similar in every layer, they might be highly dissimilar in every layer, or they might begin similar to each other, but diverge over the course of processing, similar to the pattern of the six example objects in Figure 1e.

**Figure 2.**
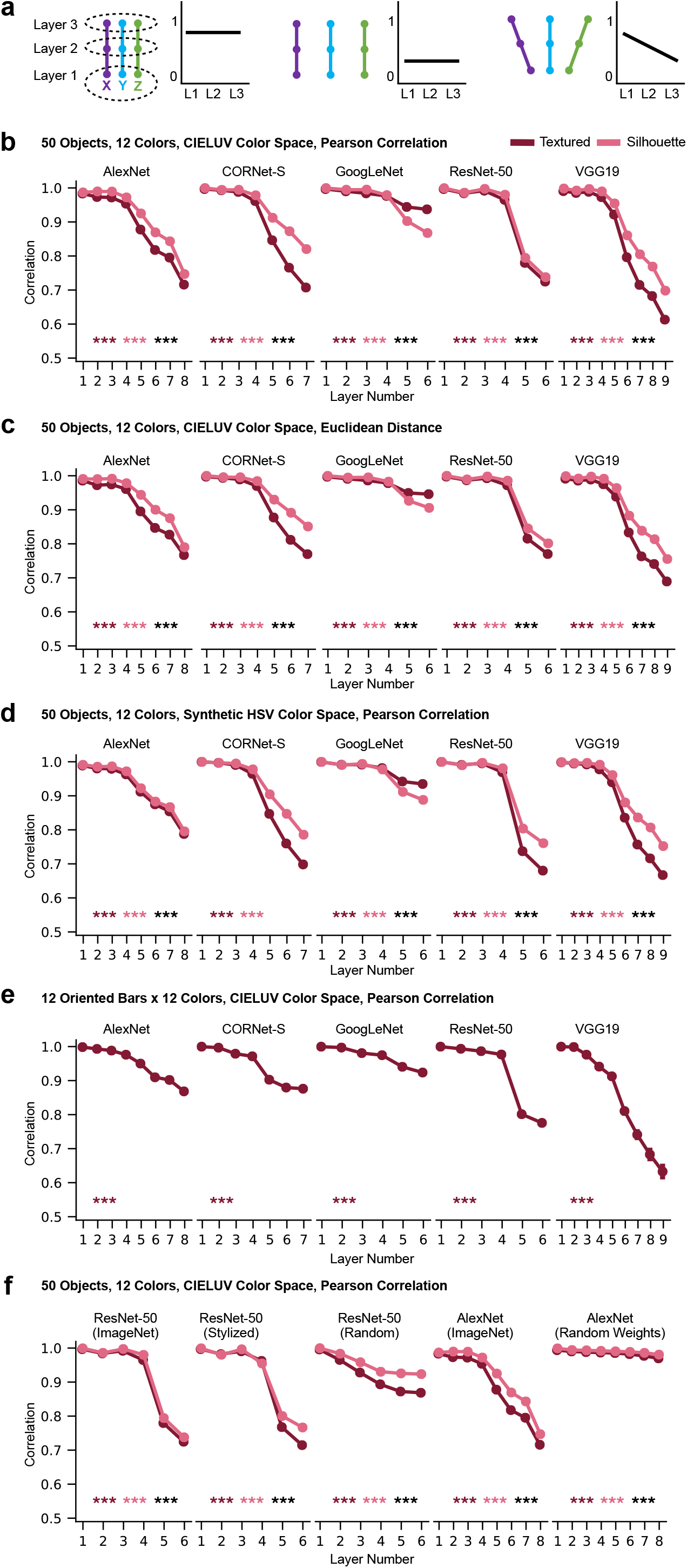
Color space representation across objects within a CNN layer. **a.** A schematic illustration of three possible scenarios. In each scenario, the left figure illustrates the color space transformation of three objects in three hypothetical CNN layers, with each colored dot depicting a color space structure of an object at a CNN layer and each trajectory depicting an object. The right figure in each scenario illustrates how the mean pairwise correlation of all object color spaces for a given layer changes across layers. In the first scenario, the color spaces of the three objects remain similar within each layer throughout processing. In the second scenario, they are dissimilar within each layer throughout processing. In the third scenario, they are similar in the first layer, but become dissimilar in the final layer. **b.** The mean pairwise color space similarity for each sampled layer of each of the five CNNs, for the full set of 50 objects in the 12 colors calibrated in CIELUV color space. The color space structure of each object is measured with Pearson correlation. Results are shown for both the textured (in maroon) and the silhouette (in pink) versions of the objects. Linear regression was used to measure the downward trend of the mean correlations across layers for each object pair. The mean of the resulting slopes (one slope per object pair) were tested against zero for each of the two versions of the objects, and the difference between the two sets of slopes was also tested (with significance levels marked by maroon and pink asterisks, respectively, for each of the two versions against zero, and by black asterisks for the differences between the two versions). In all cases, mean pairwise color space similarity decreases over the course of processing, with this decline being greater for the textured than for the uniformly colored objects. **c.** Same as (**b**), but with color space structure of each object measured with Euclidean distance instead of Pearson correlation. Results remain qualitatively similar as those in (**b**). **d.** Same as (**b**), but using colors calibrated in an artificial HSV color space that is not based on human psychophysical judgments. Results again remain qualitatively similar as those in (**b**). **e.** Mean color space similarity across the 12 oriented bar stimuli in the 12 colors calibrated in CIELUV color space. Even in these minimally simple shape stimuli, mean pairwise color space similarity decreases over the course of processing. **f.** Mean color space similarity across the 50 objects in the 12 colors calibrated in CIELUV color space in CNNs with different training regimes. Comparisons are made among ResNet-50 trained with the original ImageNet images, trained with stylized ImageNet images, and with 100 random-weight initializations of the network. Comparisons are also made between AlexNet trained with the original ImageNet images, and with 100 random-weight initializations of the network. Averaged results are shown from the 100 random-weight initializations of each network. The untrained networks exhibit a much smaller decline in their mean pairwise colorspace correlation across objects than the trained networks. † p<.1, * p<.05, ** p<.01, *** p<.001.

Figure 2b depicts the mean between-object color space correlation within each layer as well as how the mean changes across layers. In all CNNs, and in both stimulus conditions (textured and silhouette), the mean color space correlations were high in lower layers but then significantly decreased from mid to high CNN layers, with an overall significantly negative slope across all the layers (see the asterisks marking the significance level of the slope at the lower part of each plot). Coding of color thus remained relatively similar across objects in lower layers but then became increasingly different from mid to high CNN layers, reflecting what is depicted in Figure 1e and Figure 2a right panel. Since this increase in interactive tuning occurred even for the silhouette stimuli, it did not depend on the internal texture features of the stimuli, and can occur with respect to global shape features alone. That being said, for most of the networks, the textured stimuli did exhibit a greater drop in their color space similarity over the course of processing than the silhouette stimuli, with the exception of GoogLeNet, suggesting the existence of greater interactive coding for texture features above and beyond global shape features alone.

To ensure that these results did not arise due to the particular similarity metric used in constructing the RSMs (i.e., Pearson correlation), we repeated the same analysis using Euclidean Distance as the distance metric (Figure 2c). Additionally, to ensure that our results did not depend on the human-based color space we used (i.e., CIELUV color space), we repeated the same analysis, but using stimuli where the saturation and luminance were equated according to a new color space we constructed, “synthetic HSV”, which was not based on human psychophysical measurements (Figure 2d). For both manipulations, the results remained qualitatively similar: color space correlations among different objects began high in early layers, and dropped significantly in later layers in all conditions.

Could these results have arisen due to the objects subtending different areas of space? Some stimuli, like the top hat, covered large areas, while other stimuli, like the necklace, covered small areas (Figure 1a). This could have activated different numbers of kernels in CNN layers and affected how colors are coded for each object. To investigate this possibility and to examine whether our results hold for minimally simple stimuli, we repeated the same analysis on 12 oriented bars presented in the same 12 colors equated in CIELUV color space as used earlier (Figure 1c). We found the same overall result: as processing proceeds in each network, color coding increasingly differs across different shape features (Figure 2e). Thus our results hold for objects equated in their spatial coverage. Moreover, results obtained from complex natural objects can be generalized to simple shape stimuli.

Overall, across all conditions we examined, we found a consistent pattern: all CNNs showed near-orthogonal color and shape processing in early layers, but increasingly interactive feature coding in higher layers.

### The Effect of Training on CNN Color Space Representation

The ImageNet images used to train the CNNs studied so far contain real-world objects with natural color-shape covariation (e.g., bananas are yellow). Could the interactive color and shape coding observed so far in CNNs be driven by such covariation in the training images? To address this question, we compared results from the version of ResNet-50 trained on the original ImageNet images and the version trained on stylized ImageNet images in which the original texture and color of every single image was replaced by the style of a randomly chosen painting, removing the real-world color-shape covariation in the natural objects (Geirhos et al., 2019). Interestingly, the version of ResNet-50 trained on stylized images still exhibited a significant, steep decrease in their color space correlation over the course of processing (Figure 2f), almost as steep as that observed in the version of ResNet-50 trained on the original ImageNet images. This suggests that the interactive color and shape coding observed in CNNs does not rely on the presence of consistent color and shape pairing naturally occurring in the training images. That said, the slopes were slightly, though significantly, steeper for the version of ResNet-50 trained on the original than the styled images of ImageNet in both stimulus conditions (*ts* > 2.89, *ps* < .004). Thus training on naturalistic images does appear to increase the degree of interactive color and shape coding in this CNN, although the effect is fairly small.

To understand the extent to which the effects we observe may arise due to the intrinsic architecture of the networks versus being a result of object classification training, we examined 100 random-weight initializations of AlexNet and 100 random-weight initializations of ResNet-50, and compared the results with those from the ImageNet image-trained AlexNet and ResNet-50 and the stylized ImageNet image-trained ResNet-50. Results for the 100 random initializations of each network were computed independently, and then averaged together. As shown in Figure 2f, while the random networks still exhibited a significant decline in their mean pairwise colorspace correlation across objects, this decline was small, and much smaller than in the corresponding trained version of each network (matched-pairs t-tests; *ps* < .001).

Overall, these results show that the intrinsic CNN architecture is not sufficient to give rise to the large interactive color and shape coding observed so far. Training on the object classification task, even with inconsistent pairings of color and shape in the object stimuli, appear to play a significant role in creating such coding.

### Transformation of Color Space Representations Across CNN Layers and Architectures

Instead of focusing on color space differences across objects within a layer, here we took an orthogonal approach and tested how the color space of a given object may change across layers by correlating the color space vector of a given object between layers. Color coding for a given object may remain similar across layers, resulting in closely clustered color spaces across layers (Figure 3a, left), or it may transform substantially over the course of processing, leading to dispersed color spaces (Figure 3a, right). To quantify such transformations, for each object, we correlated the color space vector from each layer with the color space vector from the first and penultimate layers of the network (Figure 3b). We then used regression to examine whether the correlation significantly decreases with an increasing number of intervening layers from the reference (first or penultimate) layer. Across the main set of 50 objects in 12 colors calibrated in CIELUV color space, in all cases, there was a significant and steady decrease in correlation with the target layer with increasing number of intervening layers (see the asterisks marking the significance level of the slope at the lower part of each plot). Even for the first few layers, although color space correlations within a layer were fairly high among the different objects (see Figure 2b), the color spaces of each object still differed *across* layers. Overall, color space was successively and substantially transformed over the course of processing, with the correlations between the color spaces at the beginning and end of processing being quite modest.

**Figure 3.**
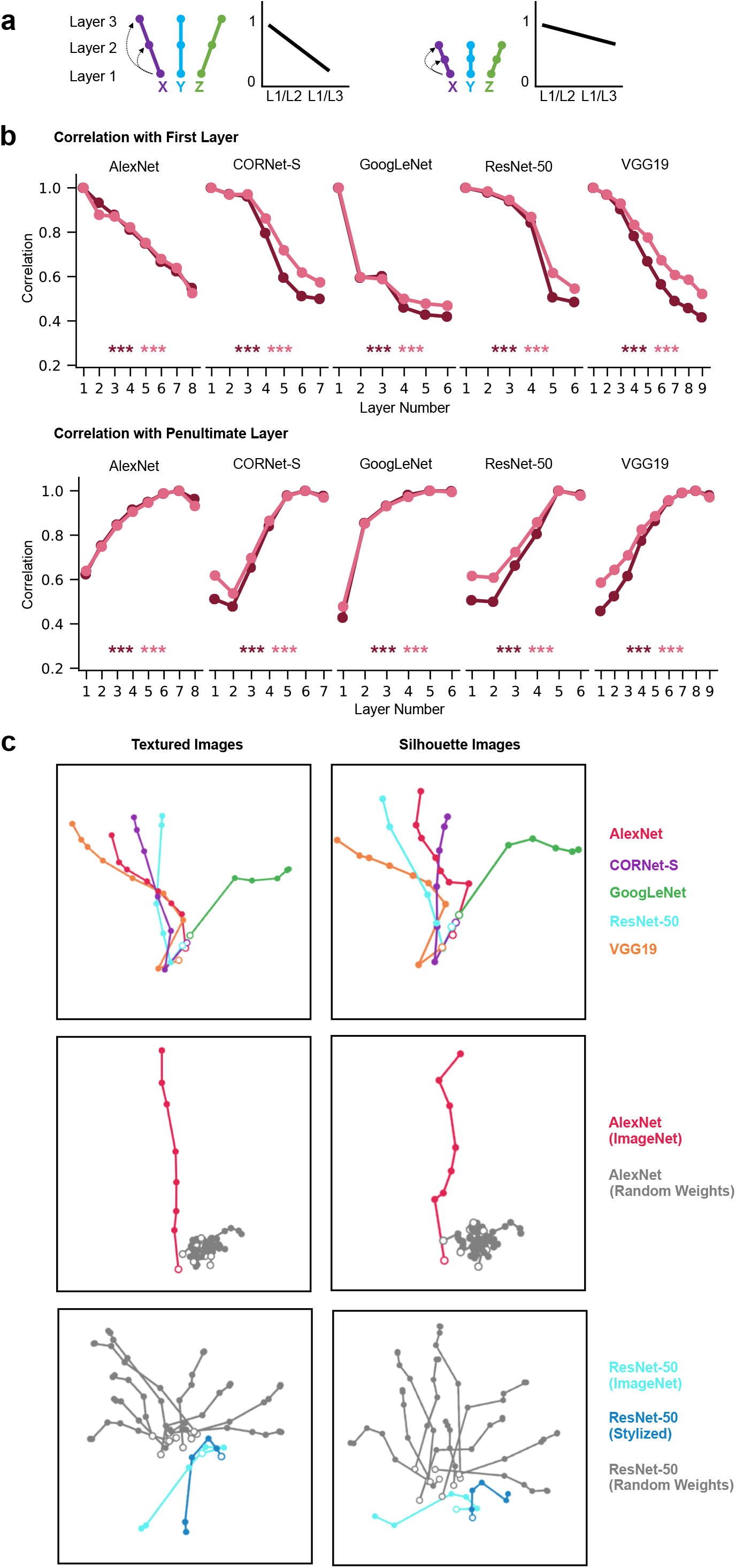
Color space representation across different CNN layers and different CNN architectures. **a.** A schematic illustration of two possible scenarios of color space correlation across layers within a CNN, using the same notations as those in **Figure 2a**. In this analysis, within each object, the color space structure from the first layer is correlated with each of the other layers, as shown on the left of each scenario. The averaged correlation overall all objects for each layer is plotted in a line graph on the right of each scenario. In the first scenario, the color space structure within each object differs substantially across processing, resulting in a large decrease in correlation across layers. In the second scenario, the color space structure for each object remains relatively stable across processing, resulting in a relatively small decrease in correlation across layers. **b.** Mean within-object across-layer color space correlations for each network for the full set of 50 objects in the 12 colors calibrated in CIELUV color space for both versions of the objects. Top row shows the correlations with the first layer of each network, bottom row shows correlations with the penultimate layer of each network. Linear regression was used to measure the downward or upward trend of the mean correlations across layers for each object. The resulting slopes were tested against zero for each of the two versions of the objects (with significance levels marked by maroon and pink asterisks, respectively). In all cases, the color space similarity within an object significantly decreases with more intervening layers, and correlations between early and late layers were fairly modest. **c.** MDS plots depicting color space correlation across different CNN layers and architectures. This was done by constructing a color space correlation matrix for each object, including its color space correlation across all sampled layers of all CNNs. The resulting correlation matrix was then averaged across objects and visualized using MDS. This was performed for the five trained networks (left column), AlexNet trained with ImageNet images and with 10 random-weight initializations (middle column), ResNet-50 trained with ImageNet images, trained with stylized ImageNet images, and with 10 random-weight initializations (right column), and for both the textured and silhouette images. The hollow dots denote the first layer of each network. In the 5 trained CNNs (left column), color spaces are almost identical in the first layer and then gradually fan out during the course of processing. Color spaces in the untrained networks, however, differ substantially from the trained ones (middle and right columns). † p<.1, * p<.05, ** p<.01, *** p<.001.

To understand how the color space of an object may be encoded differently among the different CNNs, for each of the 50 objects, we also correlated its color space vector across all CNNs and layers. We then visualized the results, averaged over all objects, using MDS plots. As shown in Figure 3c, across the 5 CNNs, for both the textured and silhouette objects, while the color spaces evolved substantially from their initial state over the course of processing (consistent with the quantitative analyses above), the color representations nonetheless evolved in a relatively similar way across networks, with the representations being almost identical in the first layer for all 5 CNNs and then gradually fanning out during the course of processing, with GoogLeNet showing a greater divergence compared to the other CNNs (see also Supplementary Figure 1 for the exact between-network correlation values for both the initial and penultimate layers of each network).

To further understand how training on object classification may affect the color space of an object, we repeated the above analysis and correlated the color space vector of the same object across the same network under different training regimes: (1) across AlexNet trained with ImageNet images and 10 random initializations of AlexNet; and (2) across ResNet-50 trained with the original ImageNet images, trained with stylized ImageNet images, and 10 random initializations of ResNet-50 (Figure 3c and Supplementary Figures 2 and 3). In both cases, while color was initially encoded in a similar manner between the random and trained versions of the networks, over the course of processing, the color spaces for objects in the trained networks substantially diverged from those in the random networks. Interestingly, while the color spaces of the different random initializations of AlexNet tended to cluster together over the course of processing and did not diverge as the trained network did, those of the different random initializations of ResNet-50 diverged substantially but in different directions as those of the trained networks. On average, in the penultimate layers, color spaces for the two trained versions of ResNet-50 tended to be more correlated with each other than they are with the random initialization of the network; this was more so for the textured than the silhouette version of the objects.

Overall, these results demonstrate that, within a given network, color representations for each object transform dynamically over the course of processing. Across the different networks, even though all the trained networks succeed at object recognition, and even though they all formed increasingly interactive representations of shape and color as processing proceeds, the exact manner in which they conjunctively coded shape and color varied somewhat from network to network, with some being more similar than others. Such transformation of color space was not a mere byproduct of a network’s intrinsic architecture, and differed substantially between the trained and untrained networks.

### Transformation of Color Space Similarity Across CNN Layers

To understand how color space similarity may change across objects over different CNN layers, instead of testing the color space of a single object, here we asked: if two objects have a relatively similar color space at the beginning or end of the network, do they also have a relatively similar color space throughout the network (Figure 4a left), or do the relative similarity of their color spaces transform throughout processing (Figure 4a right)? To test this, for the main set of 50 objects in the 12 colors calibrated in CIELUV color space, for each CNN, we took either the first or penultimate layer as our target layer and first generated its color space similarity RSM by performing all pairwise correlations of color space vectors between objects. We then vectorized the off-diagonal elements of this RSM to form a “color space similarity” vector and correlated this vector with those from all other layers of the CNN. We found that correlations decreased as we moved away from the reference layer, with the first and last layers being only moderately correlated. Thus, if two objects had a highly similar color space at the beginning of a CNN, they did not necessarily have a highly similar color space at the end of the CNN. Color space differences among objects appear to dynamically change throughout the course of processing.

**Figure 4.**
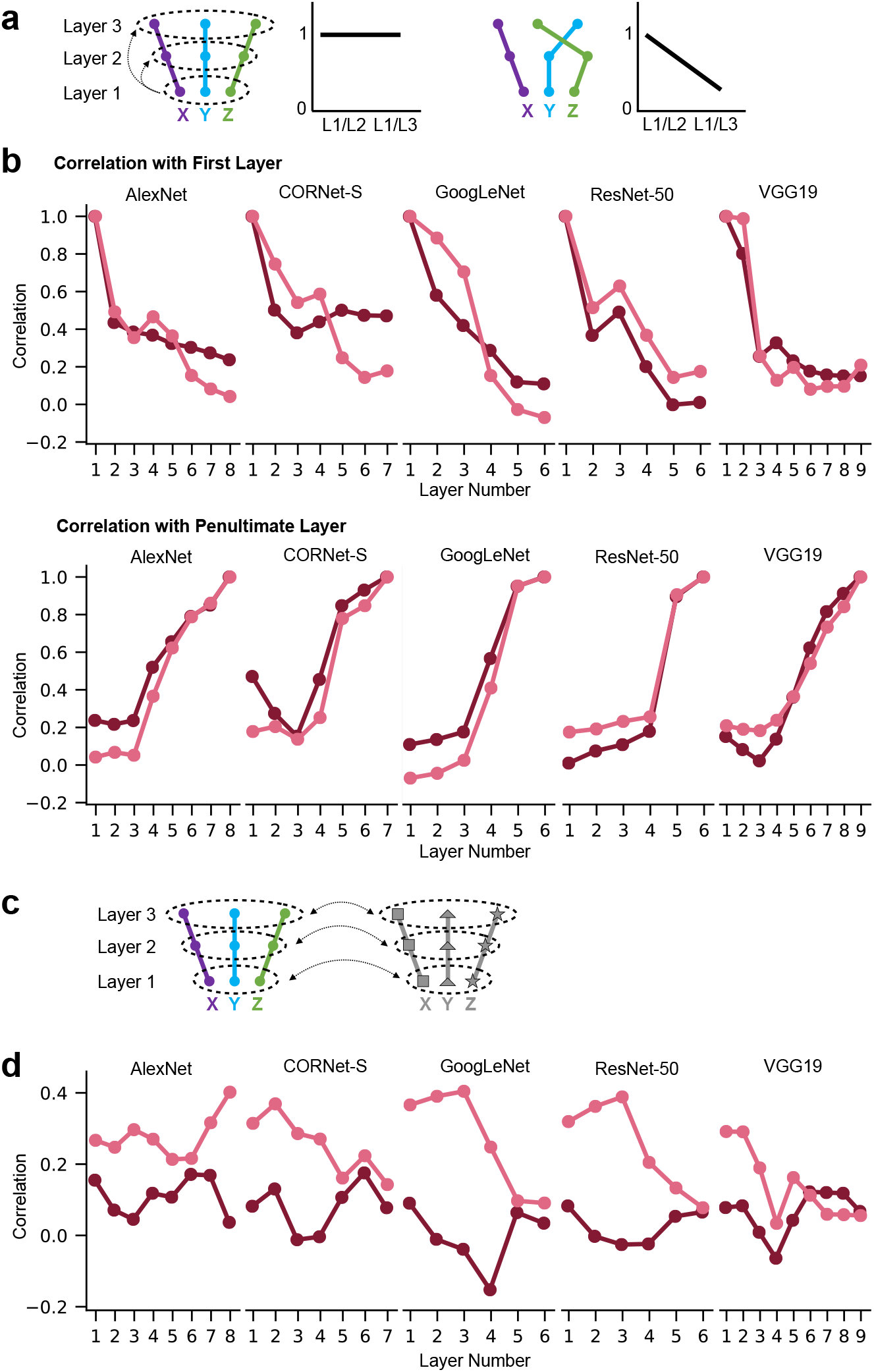
The evolution of color space similarity among objects across CNN layers and the dependence of color space similarity on object shape similarity. **a.** A schematic illustration of two possible scenarios of the evolution of color space similarity among objects across CNN layers, using the same notations as those in **Figure 2a**. In this analysis, we examine whether or not color space similarity among objects are preserved across layers by correlating the color space similarity matrix (i.e., the second-order RSM quantifying the similarity among the color spaces of different objects) from the first layer with each of the other layers, as shown on the left of each scenario. These correlations are then plotted in a line graph on the right of each scenario. In the first scenario, the relative color space similarity among the different objects is preserved in the different CNN layers (i.e., the purple dot is closer to the blue than the green dot across the whole network), even as the absolute similarity among color spaces decreases. In the second scenario, the relative color space similarity is not preserved in different CNN layers (i.e., the purple dot is closer to the blue than the green dot in the first layer, but is closer to the green than the blue dot in the last layer). **b.** The correlations of the color space similarity across different CNN layers for the full set of 50 objects in the 12 colors calibrated in CIELUV color space for both versions of the objects. Top row shows the correlations with the first layer of each network, bottom row shows correlations with the penultimate layer of each network. In most cases, correlations between the early and late layers are fairly modest. **c.** A schematic illustration of comparing color space similarity and object shape similarity, using the same notations as those in **Figure 2a**. In this analysis, the achromatic object shape similarity matrix is extracted for each CNN layer and then correlated with the corresponding color space similarity matrix of that layer. **d.** Correlations between the shape similarity and color space similarity for each layer of each network for the full set of 50 objects in the 12 colors calibrated in CIELUV color space for both versions of the objects. No clear increases or decreases across layers were evident, but in general correlations were modest, never exceeding .4.

### The Effect of Object Shape Similarity on Color Space Similarity

It is possible that color space similarity covaries with object shape similarity, such that a small change in shape features leads to a small change in the associated representational geometry for color. However, given that each feature can vary relatively independently of the other feature, it is also possible that color coding does not closely follow shape coding. To arbitrate between these two possibilities, for the main set of 50 objects in the 12 colors calibrated in CIELUV color space, for each CNN layer, we first performed all possible pairwise correlations of the CNN layer output for the grayscale versions of the object shapes to form an object shape similarity RSM and vectorized the off-diagonal elements of this RSM to form an object shape similarity vector. We then correlated this object shape similarity vector with the color space similarity vector from the same CNN layer (Figure 4c). If similar object shapes had similar color space structure, we expected to obtain a high correlation between the two. As shown in Figure 4d, correlations varied across layers and networks, showing no consistent pattern; correlations tended to be higher for the silhouette than for the textured stimuli, but tended to be modest, never exceeding a value of *r* = 0.4. Overall, color space similarity does not closely track object shape similarity, suggesting some separation between the two.

## Discussion

Despite decades of neuroscience research, we still lack a full understanding on how feature conjunctions are represented in the primate brain. In this study, we took advantage of the recent development in CNNs trained to perform object classification and examined how such an information processing system jointly represents different object features across the entire processing hierarchy of a CNN. Our investigation not only allowed us to gain insight into the internal representations of CNNs, but also enabled us to develop a novel network-based stimulus-rich approach to study feature binding across the entire network and a large stimulus set, which can be easily implemented to study feature binding in biological visual systems. Although we tested the joint coding of color and shape here, our approach can be applied to study the joint coding of any pair of features.

With this approach, we found that color coding increasingly varies across different real-world object shapes in higher levels of each CNN. This held true for both the naturally textured stimuli and the uniformly colored “silhouette” stimuli, suggesting that interactive coding of color and shape in higher CNN layers exists for global shape features alone (which are preserved in the silhouettes). The textured interior of an object shape, however, did further increase the amount of interactive coding between colors and shapes, likely due to the presence of additional shape features in these textured objects. This interactive coding was present not only for complex real-world object shapes, but also for minimally simple oriented bar features when the size of the stimuli are equated; in other words, color coding differed greatly even among otherwise identical bars of different orientations. Finally, this effect did not depend on the distance metric that was used to compute the representational geometry of the features, nor did it depend on our particular selection of color space used to calibrate the luminance and saturation of the images.

The interaction between color and shape coding was not a mere byproduct of the intrinsic architecture of a CNN, as the interaction effect was profoundly attenuated in untrained CNNs with random weights. Neither did it appear to depend on the existence of natural covariation between shape and color in the training set, as the magnitude of interactive tuning is nearly as large in a CNN trained on objects stripped of their naturalistic shape-color pairings. Thus training for object recognition is needed to produce the interactive coding of color and shape, even when no consistent color and shape pairing is present during training. This suggests that the interactive coding of color and shape is not intrinsically tied to the CNN architecture and that object recognition training automatically gives rise to increasingly tangled color and shape representations in higher levels of processing, even when color is not informative to object recognition after training.

In additional analyses, we found that an object’s color space greatly diverged from its initial color space over the course of processing and that two objects with a similar color space at the beginning of processing did not necessarily have a similar color space at the end of processing. Thus the color space representation for a given object as well as the relative similarity of color spaces between objects dynamically changed over the course of processing. However, the color space of an object tended to transform in similar ways across the trained networks, but differently in the untrained networks. This relative consistency across the trained, but not the untrained, networks with vastly varying architectures suggests that this resculpting of color space may be of adaptive value for the network’s object classification task. Interestingly, the achromatic shape similarity of two objects only weakly predicted the similarity of their respective color spaces. This demonstrates that, in general, color space similarity does not closely track object shape similarity, suggesting some separation between the two.

Overall, these results show that colors are not represented similarly across different objects in an orthogonal manner in CNNs. Rather, colors are encoded increasingly differently across objects in an interactive and object-specific manner during the course of CNN processing. This is more consistent with a late integration account (but without needing an additional binding operation), rather than color and shape being represented in an initially entangled and intermingled fashion and only being represented separately and explicitly in later layers. To our knowledge, these results provide the first detailed and comprehensive documentation of how color and shape may be jointly coded in CNNs, unveiling important details regarding the inner workings of CNN, which up to now have remained largely hidden.

It should be noted that interactive tuning does not imply that there exist units tuned exclusively to a single color/shape conjunction (a “grandmother unit”); units could plausibly be tuned to heterogeneous combinations of color and shape combinations in a “mixed selectivity” coding scheme. Such a coding scheme has been reported in the macaque prefrontal cortex for the coding of stimulus identity and task and has been shown to vastly increase the neural representational capacity of that brain region (Rigotti et al., 2013). The present results show that such an interactive coding scheme may be more prevalent and can automatically emerge in a complex information processing system even though, compared to a biological brain, CNNs have a relatively simple structure, consisting only of a single feed-forward sweep and lacking mechanisms such as feedback connections (except for the recurrent network we included here) and oscillatory synchrony. Such a coding scheme may well be used by sensory regions in the primate brain to support the flexible encoding of a wide range of sensory feature combinations. Indeed, although initial evidence from visual search (Treisman and Gelade, 1980) and neuropsychology studies (Zeki, 1990) suggests that color and shape might be initially encoded independently, and only combined in a late binding operation, other strands of evidence suggest that color and shape might be encoded in an interactive manner early on during processing (Rentzeperis et al., 2014). For example, Seymour et al. (2009) found that nonlinear tuning for color/orientation combinations might emerge as early as V1, V2, V3, and V4. Consistent with the present observation, interactive coding of color and shape has also recently been observed in the color selective neurons of macaque color patches (Chang et al., 2017).

Despite its significance in visual cognition, how feature conjunctions are coded in the human brain remains unresolved. A population code that instantiates interactive tuning for feature combinations, as we observe here, is a candidate mechanism that should be explored in more detail, and analogous analyses should be applied in monkey neurophysiology and human fMRI studies to see if similar response profiles exist in the primate brain.

To summarize, despite the success of CNNs in object recognition tasks, presently we know very little about how visual information is processed in these systems. The present study provides the first detailed and comprehensive documentation of how color and shape may be jointly coded in CNNs. Our development of a novel network-based stimulus-rich approach to study feature binding in CNNs can be easily implemented to study neural mechanisms supporting feature binding in the primate brain. Equally importantly, the discovery of the “mixed selectivity” coding scheme used by CNNs to code feature conjunctions could be a viable coding scheme that the primate brain may employ to solve the binding problem.

## Methods

### CNN Selection

We chose five CNNs in our analyses: AlexNet, CORNet-S, GoogLeNet, ResNet-50, and VGG19. These CNNs were selected based on several different criteria. AlexNet (Krizhevsky, Sutskever, & Hinton, 2012) was included for its high object recognition performance, relative simplicity, and prevalence in the literature. VGG19 (Simonyan & Zisserman, 2015), GoogLeNet (Szegedy et al., 2015), and ResNet-50 (He, Zhang, Ren, & Sun, 2015) were chosen based on their high object recognition performance and architectural diversity. Additionally, both AlexNet and VGG19 have a shallower network structure, whereas GoogLeNet and ResNet-50 have a deeper network structure. Finally, CORNet-S (Kubilius et al., 2018), a shallow recurrent CNN designed to approximate the structure of the primate ventral visual pathway, was included for its high correlation with neural and behavioral metrics. This CNN has recently been argued to be the current best model of the primate ventral visual regions (Kar et al., 2019). For most analyses, we used pre-trained implementations of these CNNs optimized for object recognition using ImageNet (Deng et al., 2009). To understand how the specific training images would impact CNN representations, we also examined responses from an alternative version of ResNet-50 that was trained on stylized ImageNet images in which the original texture of every single image was replaced with the style of a randomly chosen painting. This biased the model towards representing holistic shape information rather than texture information (Geirhos et al., 2019). Finally, in order to determine to what extent the architectural parameters of a network (number of layers, kernel size, etc.), independent of any training, affects the results, we also examined multiple initializations of AlexNet and ResNet-50 with randomly assigned weights and no training. The PyTorch implementations of all models were used, and custom scripts designed to interface with PyTorch were used for all analyses (Paszke et al., 2017).

Since these CNNs have varying, and often large, numbers of layers, we performed analyses over a subset of layers in each model, in order to simplify analysis and roughly equate the number of layers analyzed in each model. The first layer, several intermediate layers, the penultimate layer (i.e., the last layer before the object category label outputs), and the final layer (i.e,. the object category label output layer) were used for each model. Selection of intermediate layers varied based on the model, but since all CNNs we examined tended to be structured into architecturally significant “segments” (e.g., VGG19 has repeated “segments” composed of alternating conv and relu layers followed by a pooling layer, CORNet-S has “segments” meant to correspond to different visual brain areas etc.), we included the intermediate layer corresponding to the beginning or end of each “segment”. The specific layers we included are listed in Table 1. For this study, we adopt the convention of labeling layers by the kind of layer, followed by the number of times that kind of layer was used up to that point in processing (e.g., the third convolutional layer is conv3).

In cases where we wished to compare coding at the beginning and end of processing in the network, the first and *penultimate* layers were used; this is because the final layer is the category output layer, and thus the penultimate layer can be seen as the last “feature-coding” layer.

In order to compare our results across different CNNs, for some analyses we coded a variable, layer_fraction, that reflects what fraction of a network’s layers have been traversed up to a given layer in the course of a CNN’s processing hierarchy (all layer types were included). For example, the first layer in a ten-layer network would have a value of .1 for this variable, and the final layer would have a value of 1.0.

### Stimulus Selection

We used a set of real-world object stimuli from Brady et al. (2013) as our main object stimuli. This stimulus set consists of images (400 x 400 jpegs) of 540 different objects, where the colored portions of these objects are all initially of the same hue (so as to facilitate manipulating the colors of the objects in a consistent way). To derive the stimuli used in our analyses, we selected a smaller subset of these objects (as detailed below), and then manipulated the color and texture of these objects.

#### Main Object Stimuli in CIELUV Colorspace

For our main stimulus set, we chose 50 objects intended to be maximally dissimilar with respect to their high-level visual features. To do this, we converted the initial 540 objects to grayscale, ran them through AlexNet, and extracted their activations from AlexNet’s penultimate (last pre-classification) layer, FC2. We then constructed a representational similarity matrix (RSM) by computing all pairwise correlations of the CNN output from layer FC2 for each object with each other, and used this RSM to select a set of 50 objects whose mean pairwise correlation was minimally low. With this procedure, the mean pairwise similarity went from *r* = .25 (min *r* = -.02, max *r* = .92) for the original set of 540 objects, to a mean pairwise similarity of *r* = .13 (min *r* = -.02, max *r* = .78). The resulting set of 50 objects are shown in Figure 1A; visual inspection confirms that the resulting object set spans a wide range of different shape and internal texture features.

We then recolorized each of the 50 objects, roughly following the procedure outlined in Chang, Bao, and Tsao (2017). This procedure guaranteed that all stimuli had the same mean luminance and saturation over the non-background portions of the image. Equating mean luminance was necessary in order to equate each image’s contrast with the background. Equating mean saturation was necessary to ensure the validity of some of our analyses; in particular, since we examine whether color coding varies based on shape, failing to equate saturation could introduce spurious results, since a relatively unsaturated image would by definition have less hue variation. Each object stimulus was colored in each of 12 different hues, evenly spaced around the colorwheel. Specifically, we converted all images to the CIELUV colorspace, which is constructed such that equal distances in the space correspond to roughly equal psychophysical differences; this was done so that we could use the same stimuli on a future study comparing our results to those of human observers. Next, we computed the mean luminance (L) of each stimulus over the non-background portion of each stimulus, and did the same for the saturation (computed as 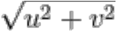, where u and v are the two chromaticity coordinates in the LUV color space). For each image, a constant value was then added to the luminance and saturation of the non-background pixels so as to bring the mean luminance and saturation of that image to target values that were equated across all objects. This procedure sometimes resulted in overflow past the permissible LUV luminance and saturation boundaries; in cases where this occurred, the variance of the luminance or saturation about the mean was shrunk until all pixel values fell within permissible boundaries. Once the luminance and saturation for each pixel were set in this manner, each object was colored in 12 different hues by rotating the U and V (hue) coordinates in each pixel to 12 equally spaced angles. Additionally, a grayscale version of each stimulus was created by setting U and V to zero, while keeping the luminance channel the same. This procedure preserves relative saturation and luminance patterns that were present across each original image, while manipulating hue and equating mean saturation and luminance. The target mean saturation and luminance were derived through successive adjustments, until applying the above transformations to each image did not take any pixel’s LUV values outside of their allowable range. Mean saturation was required to be relatively low, due to the nonlinearities of the LUV color space; specifically, a high saturation value may be possible for one luminance/hue combination, but not others, so saturation had to be kept within relatively narrow boundaries. Stimuli were converted from LUV back to RGB color space prior to being run through the networks, as the networks were trained on RGB images. Since internal texture is preserved for these stimuli, we henceforth refer to them as the “Textured” stimuli.

In addition to the above method, which preserved the internal texture of the stimuli, we also constructed a version of each stimulus that consisted of a uniformly-colored “silhouette” of the image (henceforth “silhouette” stimuli), thereby removing internal texture information while sparing global shape information. Twelve such silhouette images were created from each object, using the same 12 hues and the same mean luminance and saturation values, as were used for the Textured stimuli. We performed this manipulation because some evidence suggests that CNNs may prioritize texture over global shape features (Geirhos et al., 2019), so employing silhouette stimuli allowed us to examine whether our results hold in the absence of texture. Example stimuli from these two methods, in the 12 possible colors, are shown in Figure 1B.

#### Main Object Stimuli in Synthetic HSV Colorspace

The CIELUV color space, while widely used, is calibrated based on human psychophysical judgments, which may not be applicable to how CNNs represent visual information. To ensure that our results do not depend on these idiosyncrasies of the CIELUV color space, we constructed a new color space that was not based on human data, but more aligned with the RGB input CNNs receive. Specifically, we used a variation of the common hue/saturation/luminance parametrization calculated over RGB values, which we call Synthetic HSV. Some such parametrizations require taking maxima and minima of the RGB channels, an operation which is not available to convolutional kernels. Thus, luminance was stipulated to be the mean of R, G, and B; this definition assigns equal weights to the three channels (unlike some HSV parametrizations, which weight the three channels differently to account for human psychophysical performance), and uses an operation (taking the mean) that is easily implemented by convolutional kernels. Intuitively, saturation reflects the dissimilarity of a color from neutral grey; thus, saturation was stipulated to be the Euclidean distance in RGB coordinates from a color to neutral grey of the same luminance. Once these values are fixed, the range of possible RGB values forms a circle in the 3D RGB space. This occurs because restricting the RGB values to have a fixed average - and therefore a fixed sum - constrains the range of possible RGB values to fall on a single plane, and further restricting the RGB values to have a fixed Euclidean distance from the neutral gray point on that plane (where R = G = B) selects a circle of RGB values on that plane. Within this circle, we stipulated that the RGB triplet with the highest R channel corresponds to a hue value of 0°. Using this definition, we constructed stimuli whose mean saturation and luminance with respect to this color space corresponded to the mean values used in the CIELUV color space, with 12 equally spaced colors. Stimulus construction was otherwise identical to the procedure for the stimuli colored based on CIELUV.

#### Oriented Bars in CIELUV Colorspace

To examine the extent to which our results may hold for stimuli equated in their spatial coverage and for simpler stimuli than the naturalistic object stimuli that were used in the study, we constructed a set of oriented bar stimuli (Figure 1C). Twelve orientations, ranging in even increments from 0° to 180°, were used, and each was uniformly colored in the same twelve isoluminant and isosatured colors that were used for the object stimuli (using CIELUV space).

### Analysis Methods

For all analyses, images were fed into each network. Next, unit activations were extracted from each sampled layer and flattened into 1D vectors in cases where the layer was 3D (e.g., if it was a convolutional layer).

#### Visualizing Color Space Representation Across Objects and CNN Layers

We used representational similarity analysis (RSA) to measure conjunctive tuning for color and object shape. Specifically, we examined the extent to which the representational structure for color changes across the different object shapes. To the extent that the representational structure of color varies across the object shape, it would provide evidence that CNNs encode color and object shape not independently, but interactively.

As an initial analysis, we visualized how the color spaces for two example objects differ at the beginning and end of a CNN. Specifically, we extracted the patterns for all 12 colors of two example objects from the first and the penultimate layers of AlexNet (which are Conv1 and FC2). Within each layer, we performed all pairwise Pearson correlation among the 24 patterns to create a representational similarity matrix (RSM, with the value for each cell being the Pearson correlation coefficient). Using multidimensional scaling (MDS), we visualized the resulting similarity space (Figure 1d).

Next, we visualized how the color spaces of six representative objects might diverge over the course of processing in AlexNet (Figure 1e). To do this, for each object and for each sampled layer of AlexNet, we first constructed a “color space” RSM by performing all pairwise Pearson correlations of the patterns associated with the 12 different colors of that object (with the value for each cell of the matrix being the Pearson correlation coefficient). We vectorized the off-diagonal value of this RSM to create a “color space” vector. Next, we performed all pairwise correlations of these “color space” vectors across objects and layers to form a “color space similarity” RSM that quantifies how similarly color is coded in different objects and layers. We then used MDS to visualize the similarity of the different color spaces across different objects and CNN layers.

Following these qualitative observations, to provide a comprehensive and quantitative description of color representation across different objects and CNN layers, we performed a series of analyses. Specifically, we quantified (1) within each layer, how color is coded differently across objects (Figure 2a), (2) within each object, how color is coded across different layers of a CNN and different CNNs (Figure 3a), and (3) whether or not color space similarity among the different objects within one layer is preserved across CNN layers (Figure 4a). We also quantified how color space similarity between two objects may be determined by their shape similarity at a given CNN layer (Figure 4c). These four analyses are described in detail below. All the analyses were performed for both the Textured and Silhouette stimuli.

#### Quantifying Color Space Differences Across Objects within a CNN Layer

To understand how color is coded across objects in each CNN layer, we first created a “color space” vector for each object in each layer of each CNN as described above for our main stimulus set of 50 objects and 12 colors calibrated in CIELUV color space. We then performed all pairwise correlations of these “color space” vectors for all the objects for a given CNN layer. We next averaged these correlation values within each layer and used a line plot to visualize how the mean colorspace correlation changes over layers of a given CNN (Figure 2). To assess statistical significance of any change across layers, the correlation values between the color spaces of each pair of objects were Fisher Z-transformed and regressed onto the position of that layer in the CNN (using the layer_fraction variable described in the CNN Selection section). The resulting slope from this regression reflects the degree to which the color space similarity for these two object shapes increases or decreases over the course of the network. A positive slope would mean that the color spaces for these two object shapes become progressively more similar over the course of processing. A one-sample t-test was used to test the slopes from all possible object shape pairs against zero to assess whether the average slope was significantly different from zero. Additionally, a matched-pairs t-test was used to assess whether the slope was significantly different between the textured and silhouette images.

We also performed the same analysis in a number of control conditions, to examine whether specific choices regarding the stimulus set and analysis method affect the results. As our first control, to test how the particular similarity measure we used may impact the results, we repeated our analysis, but used Euclidean distance as our initial similarity metric instead of Pearson correlation (Figure 2c); this was done because Euclidean distance, unlike Pearson correlation, is an unbounded metric, and we sought to ensure that the choice of the specific similarity measure did not affect the results. As our second control, to examine whether our results depended on our particular choice of color space, we repeated the same analysis on the same set of objects whose colors were calibrated in the synthetic HSV space described above, instead of the LUV space used in our main stimulus set (Figure 2d). As a final control, we repeated the same analysis on simple oriented bar stimuli, where the different object “shapes” were simply different orientations of a centrally placed bar stimulus (Figure 2e). This allowed us to examine whether the results would hold for stimuli equated in their overall spatial coverage and whether results obtained from complex nature objects hold for simple shape stimuli.

#### The Effect of Training on CNN Color Space Representation

In order to assess whether the naturally occurring consistent color and shape conjunctions present in the training images were necessary to produce the results we observed, we compared models trained on naturally textured stimuli, versus unnaturally textured “stylized” stimuli (Geirhos et al., 2019). Specifically, we compared performance between ResNet-50 trained on ImageNet, and ResNet-50 trained on stylized images. In order to assess the extent to which the results are driven by the intrinsic architecture of the networks, versus being a consequence of training them for object recognition, we also performed this same analysis on 100 initializations of AlexNet and ResNet-50 with random weights and no training of any kind. The same analysis pipeline was applied to each random initialization independently, and the final results (mean color space correlation across objects within a layer) were averaged to obtain the final result (Figure 2f). Targeted matched-pairs t-tests were used to compare the slopes of these differently-trained networks with the corresponding networks trained on object recognition.

#### Transformation of Color Space Representations Across CNN Layers and Architectures

To understand how the color space of an object may evolve over the course of processing and whether colors are coded similarly for an object across different layers of a network, for the main original set of 50 objects and 12 colors calibrated in CIELUV color space, we correlated the color space vector for each object in either the first or penultimate layer of each network with its color space vector from each other layer of the network (Figure 3a). These correlation values were then averaged across all objects and plotted in Figure 3b. To test for statistical significance, we performed a regression analysis to examine whether correlations with the first and penultimate layers of the network decrease in layers that are further apart. To do this, for each object, we applied Fisher’s Z transformation to the correlation values, and regressed them onto the positions of the CNN layers (using layer_fraction) of all layers up to, but not including, the comparison layer (first or penultimate layer). A one-sample t-test was then used to test the slopes from all the objects against zero to assess whether the average slope was significantly different from zero.

To examine whether color coding within an object transforms in a similar manner across different networks, for each object we correlated its color space vector from each sampled layer of each network with every other layer (Figure 3c, left column). The resulting similarity space was visualized using MDS, and the mean pairwise similarities in the first and penultimate layers of each network were reported in Supplementary Figure 1.

To examine whether the color space of an object evolves in a similar way in trained networks and in randomly initialized networks, we performed the same analysis as described above comparing the version of AlexNet trained on object recognition with 10 random initializations of AlexNet (Figure 3c, middle column). We also performed the same analysis for ResNet, including the ImageNet-trained version, the version trained on stylized images, and ten random initializations (Figure 3c, right column). The mean pairwise similarities in the first and penultimate layers for these comparisons were reported in Supplementary Figures 2 and 3.

#### Transformation of Color Space Similarity Across CNN Layers

To understand how color space similarities across objects would change over the course of processing and whether two objects with a relatively similar color space at the beginning of processing would also have a relatively similar color space at the end of processing, for the main original set of 50 objects and 12 colors calibrated in CIELUV color space, we correlated the color space vector of all objects within each layer together to construct a color space similarity RSM for that layer. From the similarity matrix formed, we used the off-diagonal values to define a *color space similarity vector*. The resulting color space similarity vector from the first and penultimate layers were then correlated with those from each of the other layers (Figure 4).

#### The Effect of Object Shape Similarity on Color Space Similarity

To understand how color space similarity of two objects is determined by the shape similarity of these two objects (Figure 4c) and if two objects with similar shapes would also have similar color spaces, for the main set of 50 objects and 12 colors calibrated in CIELUV color space, we first measured the overall object shape similarity in each CNN layer for the original set of 50 objects. This was done by calculating all the pairwise Pearson correlations of the CNN layer output to grayscale versions of all the object shapes. From the similarity matrix formed, we used the off-diagonal values to define an *object shape similarity vector*. We then correlated the object shape similarity vector with the corresponding color space similarity vector for that CNN layer. The resulting correlation value from each CNN layer was plotted together in a line graph (Figure 4d).

## Acknowledgements

This research was supported by a National Science Foundation Graduate Research Fellowship (DGE1745303) to J.T. and a National Institute of Health Grant (1R01EY030854) to Y.X. We thank Talia Konkle, George Alvarez and other members of the Harvard Vision Lab for insightful discussion of this study.

## Author contributions

J.T. and Y.X. conceptualized the study; J.T. constructed the stimuli and performed all the analyses; J.T. drafted the manuscript with edits and comments provided by Y.X.

---

**Supplementary Figure 1.**
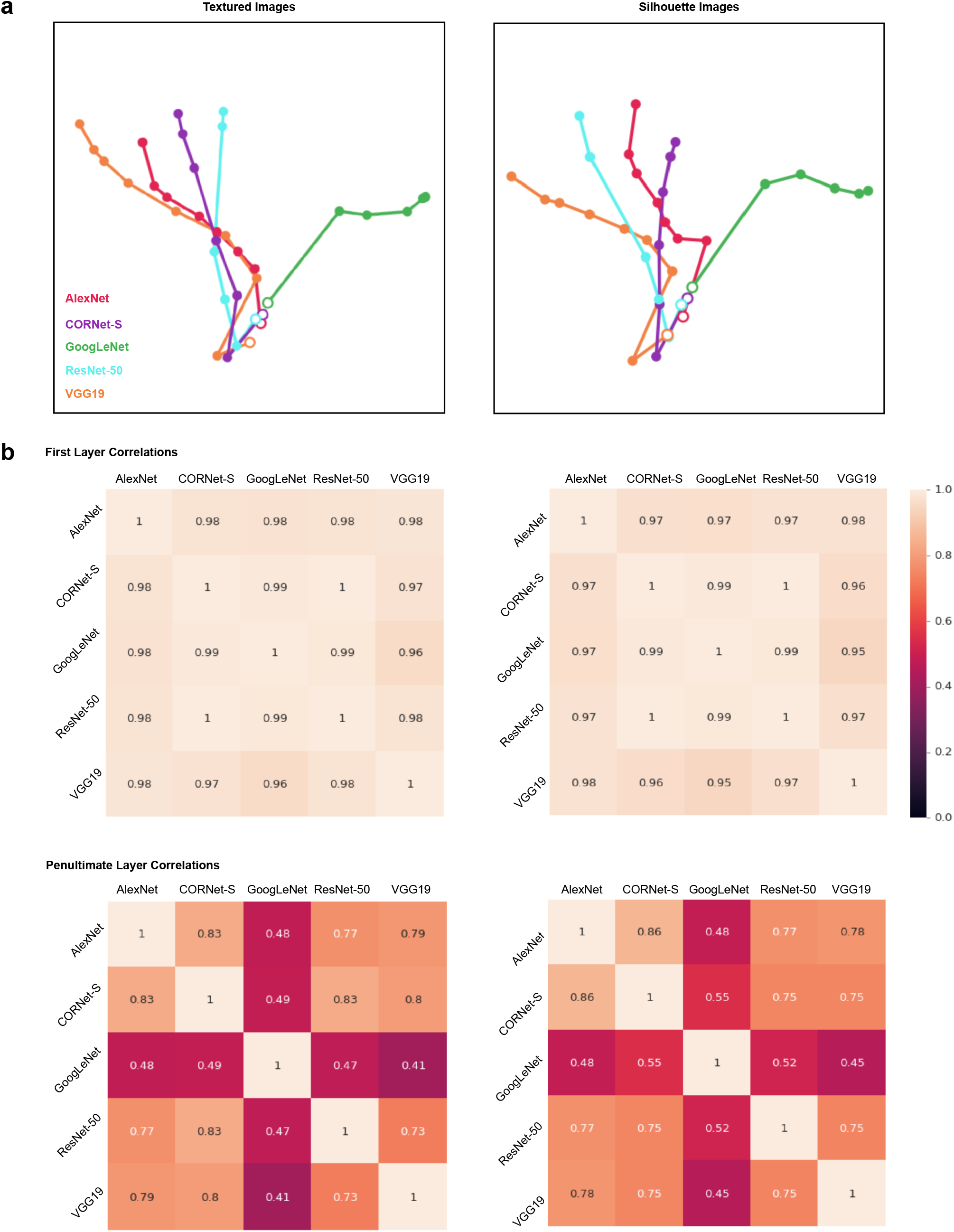
Color space comparisons for the five trained networks. **a.** MDS plots depicting color space correlation across different CNN layers and architectures, plotted separately for the two versions of the objects (copied from **Figure 3c** for convenience). This was done by constructing a color space correlation matrix for each object including its color space correlation across all sampled layers of all CNNs. The resulting correlation matrix was then averaged across objects and visualized using MDS. Each trajectory is a different network, each dot is a different layer, and the hollow dots denote the first layer of each network. **b.** Exact correlation values for the between network correlations in the first layer of each network (top) and the penultimate layer of each network (bottom) for both versions of the objects. Correlations are very similar across networks in the first layer, but diverges by the end of processing.

**Supplementary Figure 2.**
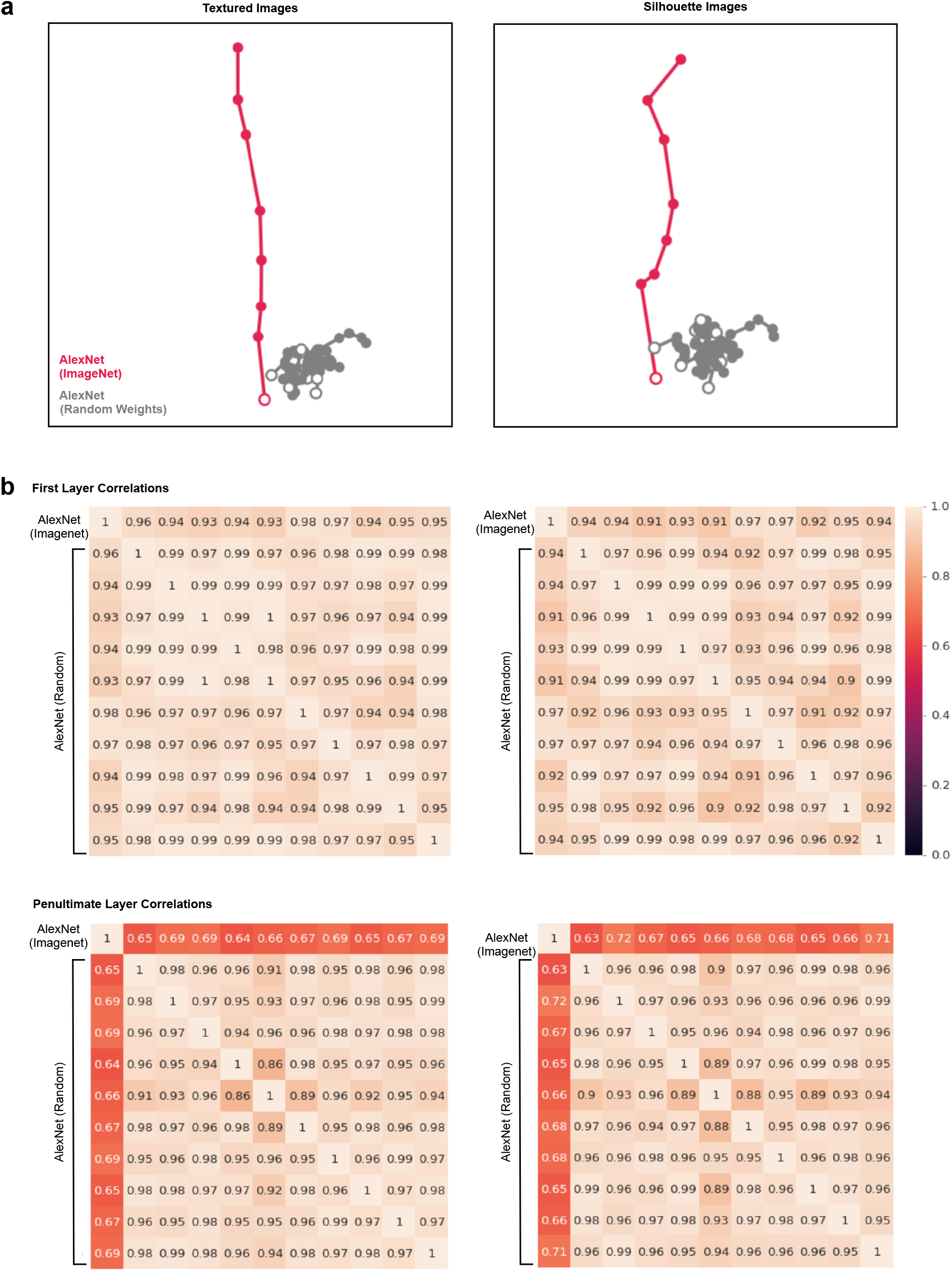
Color space comparisons for AlexNet trained with ImageNet images and with 10 different random-weight initializations. **a.** MDS plots depicting color space correlation across different layers for ImageNet trained AlexNet and 10 instances of AlexNet with random-weight initializations, plotted separately for the two versions of the objects (copied from **Figure 3c** for convenience). **b.** Exact correlation values for the between network correlations in the first layer of each network (top) and the penultimate layer of each network (bottom) for both ImageNet image trained AlexNet and the 10 instances of AlexNet with random-weight initializations, done separately for the two versions of the objects. Other details are the same as **Supplementary Figure 1**. Correlations are very similar across the trained and untrained networks in the first layer and remain similar across all the untrained networks, but differ substantially between the trained and untrained networks by the end of processing.

**Supplementary Figure 3.**
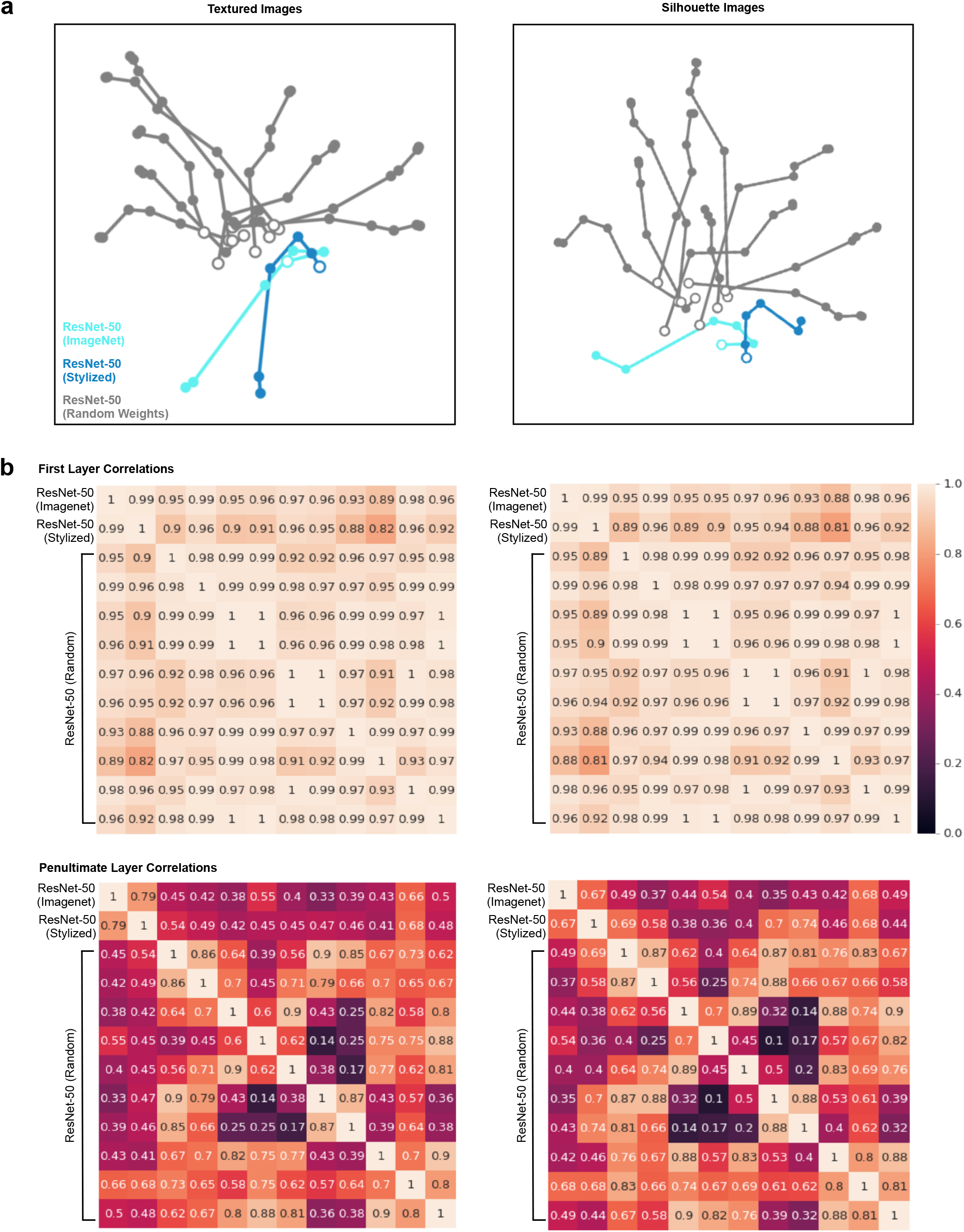
Color space comparisons for ResNet-50 trained with ImageNet images, trained with stylized ImageNet images, and with 10 random-weight initializations. **a.** MDS plots depicting color space correlation across different layers for original ImageNet trained ResNet-50, stylized ImageNet trained ResNet-50, and 10 instances of ResNet-50 with random-weight initializations, plotted separately for the two versions of the objects (copied from **Figure 3c** for convenience). **b.** Exact correlation values for the between network correlations in the first layer of each network (top) and the penultimate layer of each network (bottom) for original ImageNet image trained ResNet-50, stylized ImageNet image trained ResNet-50 and the 10 instances of ResNet-50 with random-weight initializations, done separately for the two versions of the objects. Other details are the same as **Supplementary Figure 1**. Correlations are relatively similar across the trained and untrained networks in the first layer, but diverge substantially between the trained and untrained networks, and between the 10 different instances of the untrained networks by the end of processing.

## References

Brady, T. F., Konkle, T., Gill, J., Oliva, A., & Alvarez, G. A. (2013). Visual Long-Term Memory Has the Same Limit on Fidelity as Visual Working Memory. Psychological Science, 24(6), 981–990. https://doi.org/10.1177/0956797612465439

Chang, L., Bao, P., & Tsao, D. Y. (2017). The representation of colored objects in macaque color patches. Nature communications, 8(1), 2064.

Deng, J., Dong, W., Socher, R., Li, L.-J., Kai Li, & Li Fei-Fei. (2009). ImageNet: A large-scale hierarchical image database. 2009 IEEE Conference on Computer Vision and Pattern Recognition, 248–255. https://doi.org/10.1109/CVPR.2009.5206848

DiCarlo, J. J., & Cox, D. D. (2007). Untangling invariant object recognition. Trends in cognitive sciences, 11(8), 333–341.

Flachot, A., & Gegenfurtner, K. R. (2018). Processing of chromatic information in a deep convolutional neural network. JOSA A, 35(4), B334–B346.

Geirhos, R., Rubisch, P., Michaelis, C., Bethge, M., Wichmann, F. A., & Brendel, W. (2019). ImageNet-trained CNNs are biased towards texture; increasing shape bias improves accuracy and robustness. ArXiv:1811.12231 [Cs, q-Bio, Stat]. Retrieved from http://arxiv.org/abs/1811.12231

He, K., Zhang, X., Ren, S., & Sun, J. (2015). Deep Residual Learning for Image Recognition. ArXiv:1512.03385[Cs]. Retrieved from http://arxiv.org/abs/1512.03385

Hong, H., Yamins, D. L., Majaj, N. J., & DiCarlo, J. J. (2016). Explicit information for category-orthogonal object properties increases along the ventral stream. Nature neuroscience, 19(4), 613.

Kriegeskorte, N., & Kievit, R. A. (2013). Representational geometry: integrating cognition, computation, and the brain. Trends in cognitive sciences, 17(8), 401–412.

Kriegeskorte, N. (2015). Deep neural networks: a new framework for modeling biological vision and brain information processing. Annual review of vision science, 1, 417–446.

Krizhevsky, A., Sutskever, I., & Hinton, G. E. (2012). ImageNet Classification with Deep Convolutional Neural Networks. In F. Pereira, C. J. C. Burges, L. Bottou, & K. Q. Weinberger (Eds.), Advances in Neural Information Processing Systems 25 (pp. 1097–1105). Retrieved from http://papers.nips.cc/paper/4824-imagenet-classification-with-deep-convolutional-neural-networks.pdf

Kubilius, J., Schrimpf, M., Nayebi, A., Bear, D., Yamins, D. L. K., & DiCarlo, J. J. (2018). CORnet: Modeling the Neural Mechanisms of Core Object Recognition. BioRxiv, 408385. https://doi.org/10.1101/408385

Rafegas, I., & Vanrell, M. (2018). Color encoding in biologically-inspired convolutional neural networks. Vision research, 151, 7–17.

Rajalingham, R., Issa, E. B., Bashivan, P., Kar, K., Schmidt, K., & DiCarlo, J. J. (2018). Large-scale, high-resolution comparison of the core visual object recognition behavior of humans, monkeys, and state-of-the-art deep artificial neural networks. Journal of Neuroscience, 38(33), 7255–7269.

Rentzeperis, I., Nikolaev, A. R., Kiper, D. C., & van Leeuwen, C. (2014). Distributed processing of color and form in the visual cortex. Frontiers in psychology, 5, 932.

Rigotti, M., Barak, O., Warden, M. R., Wang, X. J., Daw, N. D., Miller, E. K., & Fusi, S. (2013). The importance of mixed selectivity in complex cognitive tasks. Nature, 497(7451), 585–590.

Serre, T. (2019). Deep learning: the good, the bad, and the ugly. Annual Review of Vision Science, 5, 399–426.

Seymour, K., Clifford, C. W., Logothetis, N. K., & Bartels, A. (2010). Coding and binding of color and form in visual cortex. Cerebral cortex, 20(8), 1946–1954.

Shadlen, M. N., & Movshon, J. A. (1999). Synchrony unbound: a critical evaluation of the temporal binding hypothesis. Neuron, 24(1), 67–77.

Simonyan, K., & Zisserman, A. (2015). Very Deep Convolutional Networks for Large-Scale Image Recognition. ArXiv:1409.1556[Cs]. Retrieved from http://arxiv.org/abs/1409.1556

Singer, W. (1999). Neuronal synchrony: a versatile code for the definition of relations?. Neuron, 24(1), 49–65.

Szegedy, C., Liu, W., Jia, Y., Sermanet, P., Reed, S., Anguelov, D., … Rabinovich, A. (2015). Going Deeper with Convolutions. Retrieved December 17, 2019, from Google Research website: https://research.google/pubs/pub43022/

Treisman, A. M., & Gelade, G. (1980). A feature-integration theory of attention. Cognitive psychology, 12(1), 97–136.

Yamins, D. L., & DiCarlo, J. J. (2016). Using goal-driven deep learning models to understand sensory cortex. Nature neuroscience, 19(3), 356.

Zeki, S. (1990). A century of cerebral achromatopsia. Brain, 113(6), 1721–1777.

